# Aberrant glycosylation reveals unexpected clinical outcomes between Luminal B and Basal high stemness index breast cancer cohorts

**DOI:** 10.1101/2024.10.14.618079

**Authors:** Mia Truter, Kevin J. Naidoo

## Abstract

Cancer stem cells facilitate tumorigenesis by hijacking normal developmental pathways, the activity of which are modulated by glycosylation. Aberrant glycosylation is a key hallmark of cancer, but little is known of its functional role within the tumorigenesis system and integrated molecular and cellular cancer landscape. The clinical and phenotypic diversity in breast cancer suggests activation of various biochemical mechanisms, each likely accompanied by unique aberrations in glycosylation. Here we investigate the mechanistic links between glycosylation and breast cancer stemness for subtypes defined by the expression hormone receptors: estrogen (ER), progesterone (PR) and human epidermal growth factor (HER2). Specifically, we consider Basal and Luminal B, the two most highly stem subtypes. These two have significanlty different patient prognoses when accounting for degree of stemness. In the case of patient samples from the Basal subtype, the high stem state is protective, while in samples studied that were identified as Luminal B subtype, the high stem state becomes a risk factor over time. By undertaking a combined machine learning and bioinformatics analysis, we show that patient prognosis varies due to activation of distinct glycosylation pathways that fit into the wider tumorigenic landscape. In the high stem Basal cohort, glycosylation of immune cell surface receptors functions to maintain stemness and facilitates an activated immune response. In comparison, aberrant mannosylation and protein trafficking promote tumorigenesis through metabolic dysregulation in the high stem Luminal B cohort. These findings suggest that glycosylation plays an integral role in tumorigenesis far more so than the important role that has been identified for specific glycans, glycoenzymes or glycogens. In a systems analysis of Basal and Luminal B subtypes, we find that their aberrant glycosylation to be specific and different from each other particular with respect to high stem cases. This opens up an avenue for personalised glycosylation-based cancer diagnostics and therapeutics discovery.

**Significance:** Aberrant glycosylation gene expression screening of Basal and Luminal B breast cancer subtypes reveals differences in their phenotypic and clinical outcomes as a function of stemness helpful for personalised diagnostic and treatment.

## Introduction

The balance between stem cell self-renewal and their terminal differentiation into specialised cells is critical to the healthy development of multicellular organisms. In contrast, the objective of cancer stem cells (CSCs) is to form constantly dividing progenitor cells that never fully mature. Interestingly, activation of developmental pathways, such as Notch, Hedgehog and Wnt, is a characteristic of both cellular processes. This is no accident, as tumorigenesis is achieved through the repurposing of embryonic developmental circuitry.^1^ In a comparison between normal mammary development and mammary tumorigenesis, multiple similarities emerge, suggesting shared underlying functionalities.^2^

Glycosylation is a mechanism through which developmental processes are controlled, and dysregulated glycosylation, although often overlooked, is a feature of cancer.^3^ Several key proteins that function in stem cell maintenance are glycosylated. The degree and nature of O-glycosylation of the Notch receptors regulate their trafficking, binding affinity, stability, and activation, thereby controlling their function.^4^ Furthermore, O-GlcNAcylation is known to control the function of the well-known pluripotency transcription factors, Sox2 and Oct4.^5^

Alterations in the glycome are responsible for immune cell receptor activation, trafficking, and signalling through modifications of glycoproteins and receptors in the extracellular environment.^6^ Glycosylation may therefore contribute to an immunosuppressive or immunostimulatory state, resulting in a pro- or anti-tumorigenic microenvironment, respectively.^7,8^ Crosstalk between immune and stem cells is necessary for establishment of the stem cell niche,^9^ thus, by mediating these reciprocal interactions, glycosylation may be directly involved in regulating cancer stemness. Interestingly, different stages of cellular pluripotency are associated with distinct glycomic features that enable varying responses to stimuli even when the transcriptomic and epigenetic landscape remains constant.^10^

Breast cancer diagnosis and treatment decisions are primarily informed by inferences from clinical examinations, hormone receptor status (ER/PR/HER2), and the proliferation marker Ki-67.^11^ The present care strategy is supported by several gene-expression based tests such as Prosigna PAM50, which stratifies patients into one of five intrinsic subtypes: Luminal A, Luminal B, HER2, Basal, and normal-like.^12,13^ Despite the significant improvement that these classification strategies have brought about, heterogeneity still exists with respect to prognosis and treatment response.^14^ Therefore, improvement of breast cancer subtyping methods is an ongoing field of research.

The expression of glycoenzyme (GE) genes that control the glycosylation process can identify cancer types, and GE expression signatures overlap with breast cancer intrinsic subtype.^15^ More recently, the degree of tumour cell dedifferentiation or stemness has been proposed as a means to classify cancers.^16,17^ The degree of stemness has been associated with metastasis and tumour aggression and consequently poor patient outcome.^18^ However, in contrast it has been linked to increased immune infiltration and a positive patient outcome.^19^ These opposing prognoses suggests that the effect of stemness on the cancer progression may be context-specific.

The Basal and Luminal B (LumB) subtypes contain the largest proportion of highly stem (mRNAsi ≥ 0.5) breast cancer samples in The Cancer Genome Atlas dataset. Consequently, we investigate the interplay between aberrant glycosylation and cancer stemness in these PAM50 subtypes. This is done using a combination of unsupervised machine learning and bioinformatics techniques to show that alterations in glycosylation are reflective of both hormonal subtype and degree of stemness. We take this approach to gain insight into the underlying biological mechanisms driving stemness and explore the role of glycosylation in context-specific tumorigenesis. These results present an opportunity to develop personalised diagnostic and treatment strategies for breast cancer.

## Materials and methods

### Data acquisition and preparation

Breast cancer RNA-seq, clinical, and Simple Nucleotide Variation data were downloaded from the TCGA (https://www.cancer.gov/tcga) using TCGABiolinks^20,21^ in R version 4.4.0.^22^ The stemness index^18^ (mRNAsi, using the SC_PCBC_stemSig function) was obtained with “TCGAanalyze_Stemness” from TCGABiolinks. All female, treatment naïve Basal (n_total_ = 189, n_HS_ = 144,n_LS_ = 45) and LumB (n_total_ = 203, n_HS_ = 125,n_LS_ = 78) samples were extracted from the database.

Ensembl IDs were converted to gene symbols using the org.Hs.eg.db library.^23^ In cases where duplicate symbols exist, the entry was chosen with the highest mean expression across samples. GE genes were obtained from GlyocoEnzOnto.^24^ Tidyverse^25^ and data.table^26^ were instrumental for performing dataset manipulation.

### Denoising Autoencoder Self Organising Map (DASOM)

We developed the unsupervised machine learning method DASOM,^27^ that makes possible class separation from complex data sets. First, the input space is transformed in the denoising autoencoder phase. At the encoder end the input (in this case glycoenzyme gene expression) data, *x*, is corrupted by introducing noise into the data using a Gaussian function with a standard deviation of 0.2. This produces the corrupted input feature, 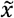. The contribution of the corrupted input feature to the hidden layer neurons are defined by weights, *W* and biasing parameters, *b*. A sigmoidal hyperbolic tangent function, *f*, is applied to these that give the activation of the hidden neurons, *y*. The size of *y* was set to 300 neurons.

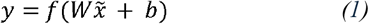

At the decoder end (the output layer) the samples’ reconstructed features, 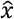, are generated. This is done via a set of parameters biasing the neurons, 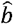, in the network’s layers accompanying the weights, 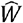. A sigmoidal hyperbolic tangent function, *f*, is once again applied to regulate of the contribution of the representation layer, *y*.

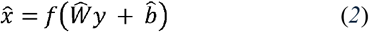

This procedure of learning features from artificially corrupted input data leads to devised representations that are close to the stable structures and invariant features present in the input data space. The DA was trained using 10000 epochs, with an initial and final learning rate of 0.7 and 0.1 respectively.

The DASOM approach is to stratify several levels of nonlinearity to the self-organizing class of algorithms. The second phase of DASOM involves training a Self-Organising Map (SOM) where the best matching neuron will coincide with the winning neuron:

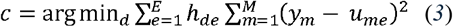

and here *E* is the number of neurons in the SOM grid, which in this case was set to 16. Every neuron *e* has an associated codebook vector *u*_*e*_ made up of *u*_*me*_ points, belonging to each individual hidden representation layer, *y*_*m*_. In these experiments the classic version of DASOM as described by us elsewhere^27^ was used, where the DA is first trained, and then the SOM and the autoencoder components are co-trained. During this phase, the initial and final SOM learning rates were set to 0.7 and 0.01 respectively, and the DA initial and final learning rates were 1.0E-7 and 1.0E-9 respectively. 10000 epochs were used in the second phase.

Here the GE genes from GlycoEnzOnto^24^ that had a standard deviation (SD) > 0.5 across the dataset (m_genes_ = 363) were used as input to DASOM,^27^ after performing variance stabilisation transformation^28^ using only the intercept term in the design matrix. Forty-five samples were randomly chosen from each group (Basal-HS, Basal-LS, LumB-LS, LumB-HS) to minimise the effect of class imbalance. A wrapper script written in R was used to run DASOM, where a brute-force grid search method was used for hyperparameterisation of the model. The purpose being to maximise the quality of clustering, which in this study was a separation of PAM50 subtypes. To assess the relationship between non-Basal subtypes, 45 samples were randomly chosen from each subtype, and 357 GE genes had a SD > 0.5. The same parameters were used as above, except that the epochs of both phases were set to 1000, and the hidden representation layer size was 357. The aweSOM^29^ and Kohonen^30^ R packages were used to visualise DASOM output.

### Differential expression analysis and candidate gene filtering

A differential expression (DE) analysis was conducted using DESEq2^28^ and log fold changes were obtained using the “apeglm” method,^31^ noting that the occurrence of a DE gene between the Basal-HS and LumB-HS groups could arise due to several reasons and not necessarily as a result of the combination of PAM50 subtype and mRNAsi.

Methodologically, we excluded genes that were a) DE between LumB-LS and Basal-LS, as that is an indication of subtype-level DE and is not a feature of high stemness, and b) DE in the same direction in both subtypes compared to their matched normal samples, since that would indicate that the gene aberration is cancer-general and not unique to the group.

Consider the following gene sets:

G_HS_ = genes DE between Basal-HS and LumB-HS with p-adj < 0.05

G_LS_ = all genes in analysis between Basal-LS and LumB-LS

LFC_LS_ = Log Fold Change of genes in G_LS_ (LFC > 0 = upregulated; LFC < 0 = downregulated)

LFC_HS_ = Log Fold Change of genes in G_HS_ (LFC > 0 = upregulated; LFC < 0 = downregulated)

G_LN_ = genes DE between LumB-HS and normal tissue with p-adj < 0.05

G_BN_ = genes DE between Basal-HS and normal tissue with p-adj < 0.05

G’_LN_ = genes not DE between LumB-HS and normal tissue with p-adj > 0.1

G’_BN_ = genes not DE between Basal-HS and normal tissue with p-adj > 0.1

The following filtering criteria was applied:

1. Identify DE genes between LumB-HS and Basal-HS G_1_ = G_HS_
2. Exclude genes DE between LumB-LS and Basal-LS in the same direction as in the HS comparison G_2_ = G_1_ □ G’_LS_ where G’_LS_ = { g □ G_LS_ | p-adj > 0.1 and LFC_LS_ ≠ LFC_HS_ }
3. Include genes upregulated in LumB-HS compared to normal tissue G_3_ = { g □ G_2_ | g □ G_LN_ } □ G’_BN_
4. Include genes upregulated in Basal-HS compared to normal tissue G_4_ = { g □ G_2_ | g □ G_BN_ } □ G’_LN_

Thus, the genes in G_3_ and G_4_ are subtype-and stemness specific and are termed candidate genes. There were 442 candidate genes, of which 19 were GEs **(Supplementary Table S1)**.

### Functional analysis

Functional analysis was conducted using the enrichGO function from ClusterProfiler^32,33^ and the Gene Ontology database.^34,35^ Basal-HS and LumB-HS candidate genes were analysed separately. Functional analysis for the upregulated genes (p-adj < 0.05, Log2FoldChange > 0.5) in the LumB-HS Late-Hazardous was also conducted with ClusterProfiler, using the MSigDB Hallmark geneset collection,^36^ which was downloaded with the msigdbr package.^37^ Rbioapi^38^ was used to predict metabolites associated with the unique differential expression (DE) signatures observed in each group. The library was set to “Metabolomics_Workbench_Metabolites_2022” (https://www.metabolomicsworkbench.org/). The four most significant metabolites from each group were chosen for visualisation, excluding alpha- and beta-D-Glucose-Phosphate in the LumB-HS group, because they were predicted from the same genes (ISYNA1 and HK1) as Glucose-6-Phosphate, which was already included in the visualisation.

### Immune cell deconvolution

The online version of Cibersort^39^ was used in absolute mode for immune cell deconvolution, utilising the LM22 signature matrix file and the TMM-normalised counts as input. Batch correction was enabled, quantile normalisation was disabled, and 1000 permutations were used for significance analysis. Only the 671 samples with a p-value < 0.05 were utilised in the analysis.

### Mutational enrichment

Mutational enrichment was performed using maftools.^40^ Basal-HS and LumB-HS mutational enrichment was performed first, and then mutations also occurring in the Basal-LS and LumB-LS comparison were excluded from the final result.

### Random forest feature discovery

The vst-normalised counts of the 19 candidate GEs was used as input to randomForest,^41^ with the objective of classifying Basal-HS and LumB-HS samples. The caret^42^ package was used to control the parameters for training (method = “cv”, k = 5, search = “grid”). Various values for ntree (100,200,500,1000) and mtry (1-30) were tried, and the model with the highest accuracy was selected. Variable importance was obtained with the varImp function.

### Other packages

Survival was analysed using casebase^43^ and the survival,^44^ GGally^45^ and survminer^46^ R packages. The alluvial plot was created with ggalluvial.^47^ Heatmaps were drawn using ComplexHeatmap.^48^ Dendsort^49^ and circlize^50^ were used for hierarchical clustering of heatmap rows and columns, and generation of colour vectors respectively. Rstatix was used for statistical testing.^51^ The Reactome^52,53^ and Genecards^54,55^ databases were frequently used to look up gene function. Transcription factors and their gene targets were accessed using The Human Transcription Factors^56^ and the TFTG databases.^57^ Oncogenes were obtained from the CancerMine database.^58^ Images were formatted for publication using BioRender.com.

## Data and code availability

All data used in this study are publicly available, and the R-code will be made available upon request.

## Statistical information

P-values were corrected for multiple testing using the Benjamini-Hochberg procedure, and significance is reported at p-adjusted < 0.05 unless otherwise stated.

## Results

### Subcluster identification of degree of stemness coupled to Basal and LumB subtypes from glycogene expression

The interplay between glycosylation, stemness, and PAM50 subtype was investigated in the two most highly stem breast cancer subtypes: Basal and Luminal B, which comprise 76% and 62% high stem samples in the TCGA (https://www.cancer.gov/tcga) dataset, respectively. High stem samples are defined as having an mRNA stemness index (mRNAsi)^18^ ≥ 0.5.

To discover whether glycoenzyme (GE) genes are able to group Basal and Luminal B breast cancer samples together in a biologically meaningful way, vst-normalised^28^ GE gene expression data (m_genes_ = 363) from Basal and LumB high (HS) and low (LS) stem samples were subjected to unsupervised clustering using the Denoising Autoencoder Self-Organising Map (DASOM).^27^ Forty-five samples were used per group to minimise the effects of class imbalance. Using partition-around-medoids superclass clustering,^29^ four clusters were found. The samples cluster largely according to a combination of PAM50 subtype and stemness index, with the dominant group in each cluster comprising around 65% of the total sample space **(Figure 1A)**.

**Figure 1:**
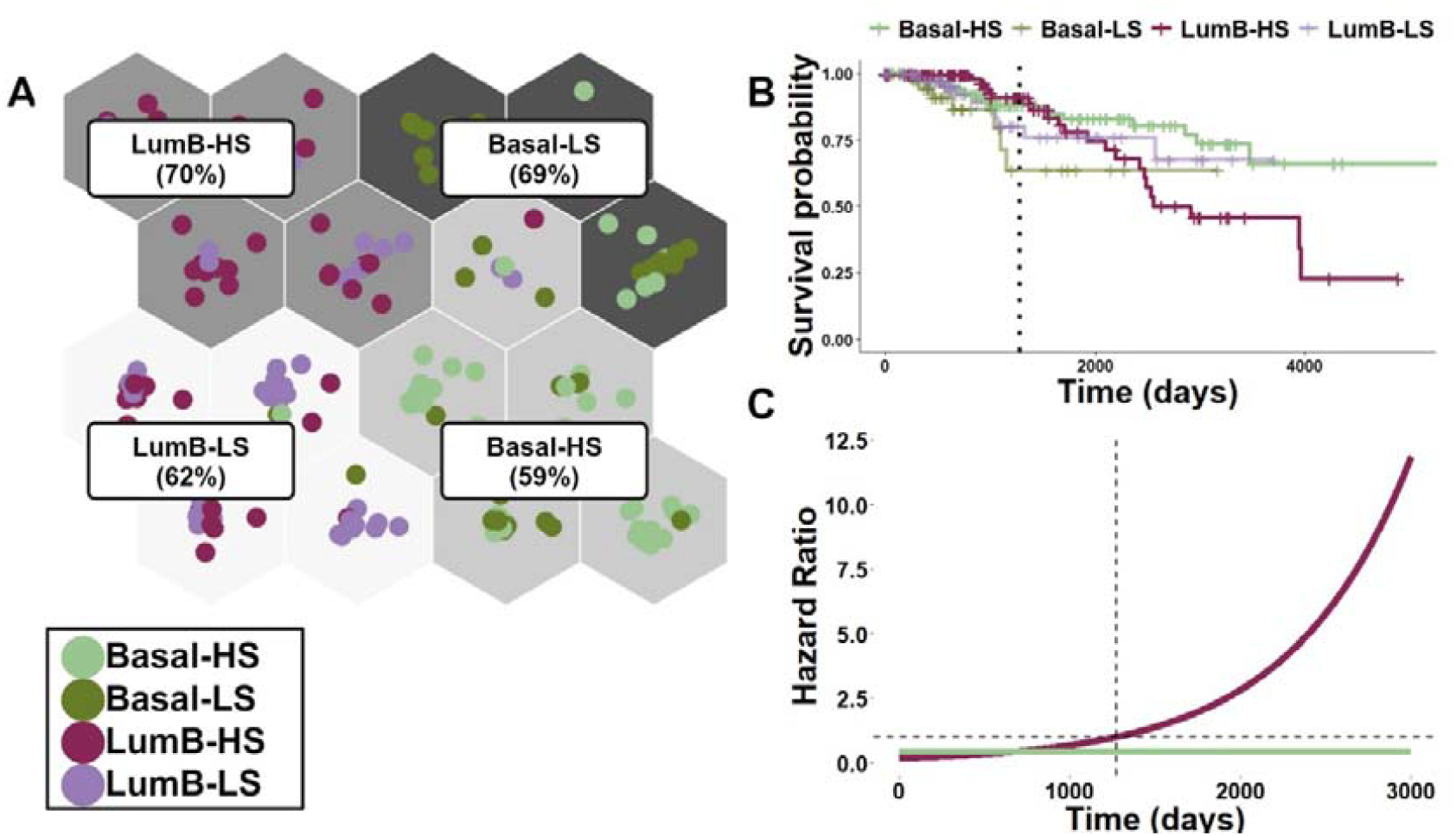
Stemness-specific subgroups within Luminal B and Basal breast cancer subtypes. **(A)** Clustering from glycogenes identify PAM50 subtype and mRNAsi signatures. The proportion of the dominant group in each cluster is shown in brackets. HS = High stem (mRNAsi >= 0.5), LS = Low stem (mRNAsi < 0.5) **(B)** Survival differences between groups. The dotted line indicates 1,273 days, where prognosis in the LumB-HS group worsens drastically. **(C)** Stemness hazard ratios over time for the LumB-HS and Basal-HS groups. Dotted line indicates 1,273 days, where the LumB-HS hazard ratio surpasses 1.

Thus, the complete Basal (n_HS_ = 144, n_LS_ = 45, n_total_ = 189) and LumB (n_HS_ = 125, n_LS_ = 78, n_total_ = 203) cohort of female, treatment-naïve TCGA samples was further investigated. The combination of PAM50 subtype and mRNAsi is clinically relevant and patient stratification based on these factors resulted in unexpected survival outcomes. While the LS groups show no significant difference in survival (Log rank p-value = 0.3), the HS groups do (Log rank p-value = 0.08) **(Figure 1B)**. Surprisingly, Basal-HS patients have a more favourable prognosis than Basal -LS patients, suggesting that stemness in the Basal subtype may be a protective factor (Cox Hazard Ratio = 0.44, p = 0.08). The stemness hazard ratio in the LumB subtype was time-dependent (Cox proportional hazards assumption p = 0.034). Modelling stemness as a categorical predictor with a time interaction^59^ showed that prior to 1,273 days, indicated by a dotted line in **Figure 1B and 1C**, stemness in the LumB group is protective, and thereafter becomes associated with an increased risk of death (hazard ratio > 1).

Limited sample size and class imbalance prevented the evaluation of the effect of metastasis and stage on patient outcome for these PAM50 subtypes coupled with the high stemness (mRNAsi ≥ 0.5) metric. The presence of the HER2 receptor was shown to be a hazardous prognostic indicator of overall survival in the LumB subtype (Cox Hazard Ratio = 2.75, p = 0.08). There was no interaction between HER2 and mRNAsi related to the overall survival in the LumB group (Cox p = 0.51) although LumB-LS samples were more likely to be HER2 -positive (chi-square p = 0.07). Pursuing this further, when LumA, LumB, and HER2 samples were subjected to unsupervised clustering, shared glycosylation signatures between LumB-LS and HER2 samples were evident **(Supplementary Figure S1)**.

### Identification of unique functional characteristics in high stem Basal and LumB samples

To understand the drivers of the differences in clinical outcomes between the LumB-HS and Basal-HS groups, a series of differential expression (DE) analyses were conducted. Genes that were exclusively DE between the LumB-HS and Basal-HS groups were obtained by filtering out genes that were DE in the LS case. Furthermore, genes that were “cancer general”, i.e., aberrantly expressed in both subtypes relative to matched normal tissue, were excluded **(Figure 2A)**. This resulted in the identification of 442 uniquely differentially expressed genes between the Basal-HS (m_genes_ = 184) and LumB-HS groups (m_genes_ = 258), termed candidate genes **(Supplementary Table S1)**. None of the candidate genes showed enrichment for Simple Nucleotide Variation mutations between groups (p-adj > 0.05).

**Figure 2:**
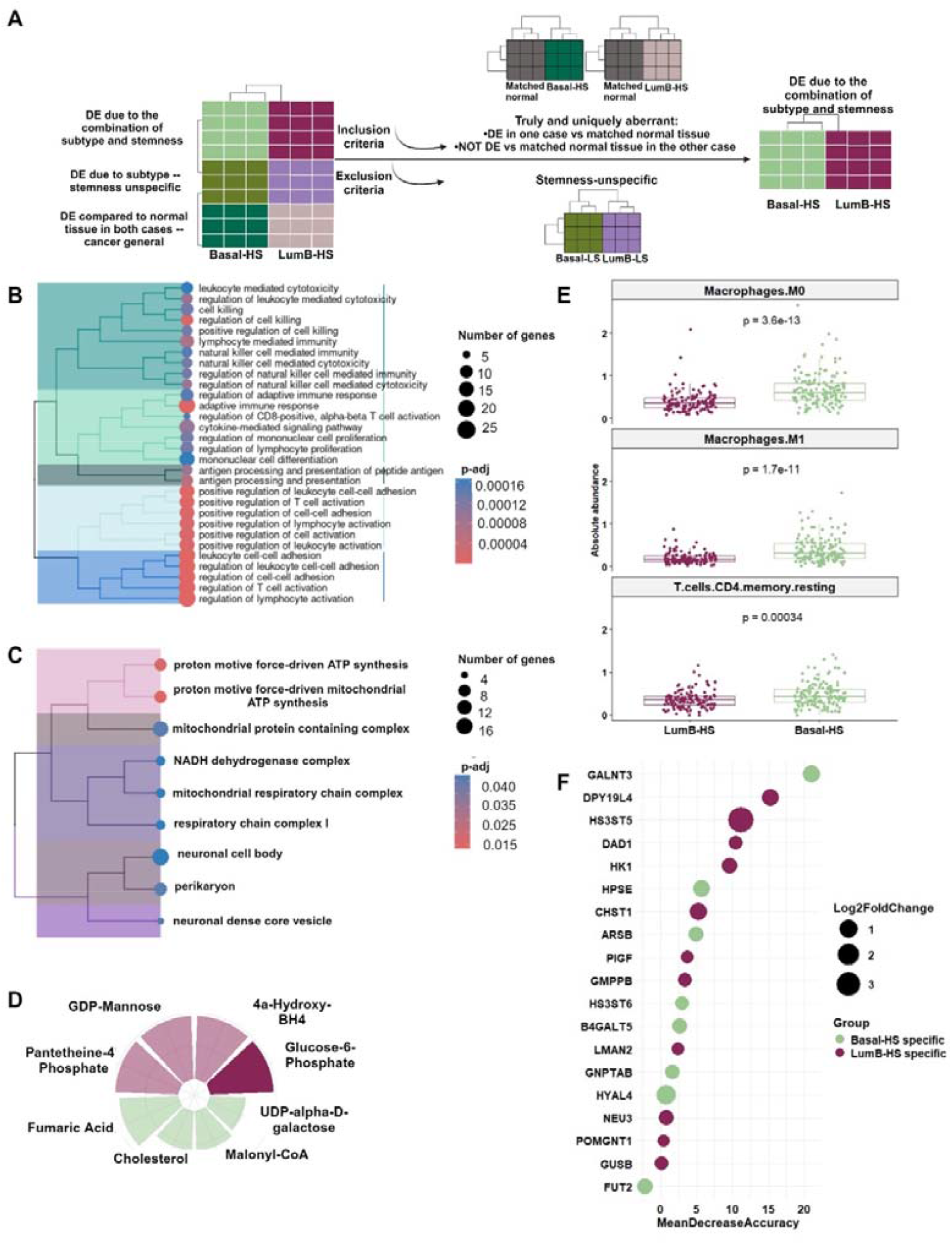
Functional characteristics of high stem Basal and LumB breast cancers. **(A)** Candidate gene identification through a series of differential expression analyses. Functional profiling of **(B)** Basal-HS and **(C)** LumB-HS groups. **(D)** Metabolite prediction of the candidate genes (hypergeometric test, p < 0.1). Size/height of wedges represents odds ratios, and translucent bars indicate metabolites that lost significance after multiple testing correction. **(E)** Differential immune cell abundance between Basal-HS and LumB-HS groups. **(F)** Variable importance of the 19 candidate GEs in classifying the Basal-HS and LumB-HS groups. Dot size indicates log fold change and dots are coloured according to which group the gene was upregulated in.

Functional profiling of the candidate genes revealed that genes in the Basal-HS group play a more active role in immune response than those in the LumB-HS group **(Figure 2B)**. Specifically, the genes that are uniquely upregulated in the Basal-HS group include programmed death ligand 1 (CD274), the major histocompatibility complexes (HLA-A/B/C), CD86 and IFNG. While the clinical implications of CD274 over-expression varies,^60^ CD274 upregulation in the Basal-HS cohort was associated with improved patient outcome (Cox Hazard Ratio = 0.73, p = 0.08). Furthermore, IFNG contributed to improved prognosis in the Basal-HS cohort (Cox Hazard Ratio = 0.63, p = 0.03). In contrast, candidate genes in the LumB-HS group included SLC45A2 and ELOVL6, which were associated with worse patient outcome (Cox Hazard Ratio = 2.49 and 2.22, respectively, p < 0.05). The expression of these genes, alongside pH-sensing ASIC3 and ATP5F1E, points to activation of the electron transport chain (aka oxidative phosphorylation) **(Figure 2C)** and suggests dysregulation of metabolic systems. Notably, the well-known stemness transcription factor SOX2 is uniquely expressed in the LumB-HS group.

To understand molecular and cellular differences between the two groups, metabolite concentrations and immune cell abundances were compared. Using hypergeometric testing, notable differences in metabolite concentrations as a result of variation in gene expression were observed **(Figure 2D and E)**. Based on the candidate genes in the Basal-HS group, UDP-alpha-D-galactose and Malonyl-CoA were predicted to be upregulated, while 4a-Hydroxy-BH4, GDP-Mannose, and Panthetheine-4’-Phosphate were enriched in the LumB-HS group **(Figure 2D)**. Furthermore, immune cell deconvolution showed that M0 (undifferentiated) and M1 (pro-inflammatory, anti-tumour)^61^ macrophages are more abundant in the Basal-HS group, as well as CD4 memory resting T-cells **(Figure 2E)**. Differential abundance of immune cell types between these groups could contribute to explaining the observed survival differences.

To determine whether the 19 candidate GEs could differentiate the two groups, they were used as input features to a Random Forest classification, and a performance of 88.47% accuracy was achieved. The most important genes contributing to the Basal-HS group classification were GALNT3, ARSB, and HPSE, HS3ST6, HYAL4, and FUT2. DPY19L4, HK1, POMGNT1 and LMAN2 played a central role in the correct classification of the LumB-HS group **(Figure 2F)**.

### Associating aberrant glycosylation with distinct biochemical pathways

While dysregulated glycogene expression is a hallmark of cancer, little is known about the functional roles that glycosylation exerts in the systems-wide tumorigenic context. The expression of candidate GEs was correlated to the remainder of the genome to uncover co-expression patterns in the LumB-HS and Basal-HS groups. Notably, genes that exhibited positive correlations to candidate GEs in the Basal-HS group were simultaneously negatively correlated to LumB-HS candidate GE expression patterns, suggesting distinct and separate GE expression patterns between these two groups **(Figure 3A)**.

**Figure 3:**
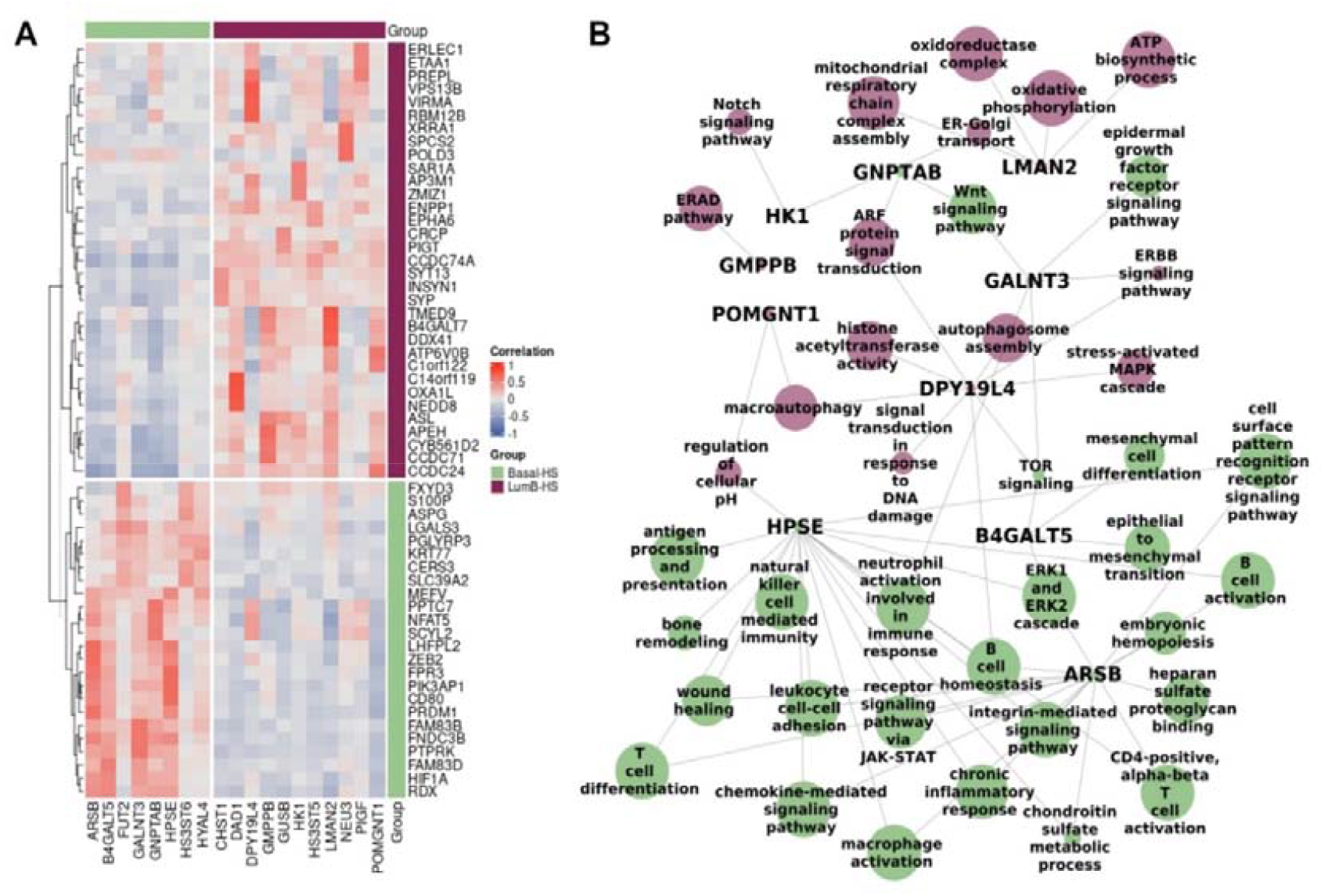
Co-expression analysis of candidate GE genes. **(A)** Candidate GEs and their top three most highly correlated genes (Pearson correlation). **(B)** Functional analysis of the genes that correlate with the candidate GEs. Only non-redundant, significant pathways (p < 0.05) are shown. Node size is inversely mapped to p-value. Nodes are coloured according to which group (LumB-HS or Basal-HS) a candidate GE is upregulated in. In the case where a pathway is shared between two candidate GEs, the node is coloured according to lowest p-value.

Genes with strong positive correlations to candidate GEs (ρ > 0.5, p < 0.05) were subjected to functional analysis with ClusterProfiler,^32,33^ revealing the manner in which GEs in each cohort may be involved in tumorigenesis **(Figure 3B)**. The LumB-HS candidate genes such as HK1, POMGNT1, and LMAN2 were correlated to genes involved in oxidative phosphorylation, responses to ER stress such as alterations of cellular pH and ER-associated degradation (ERAD), Notch signalling, and autophagy. In contrast, Basal-HS candidate GEs were associated with activation of immune cells, integrin-mediated and Wnt signalling, and epithelial to mesenchymal transition (EMT). These genes included HPSE, ARSB, B4GALT5, and GALNT3. Thus, in addition to GEs being differentially expressed between the two groups, they are correlated to genesets with distinct functions, suggesting unique and context-specific roles for glycosylation in promoting tumorigenesis.

### Time-dependent patient outcome analysis of upregulated of glyco- and oncogenes

The trajectory of the effect of high stemness on patient outcome in the LumB-HS cohort is not linearly predictable over time. After 1,273 days from the initial diagnosis, the risk of death in the LumB-HS cohort increases significantly **(Figure 4A)**. To identify the gene- and pathway-level aberrations that contribute to the increased risk of death in the LumB-HS cohort, patients were divided into two groups related to the risk of high stemness at time of death or censorship. The first group comprised LumB-HS patients who died or were censored prior to 1,025 days from initial diagnosis, which corresponds to a hazard ratio for high stemness of 0.7 (LumB-HS Early-Protective, n_censored_ = 68, n_died_ = 6). The second group of LumB-HS patients died or were censored after 1,467 days from time of initial diagnosis, when the Hazard Ratio for high stemness was 1.3 (LumB-HS Late-Hazardous, n_censored_ = 21, n_died_ = 14).

**Figure 4:**
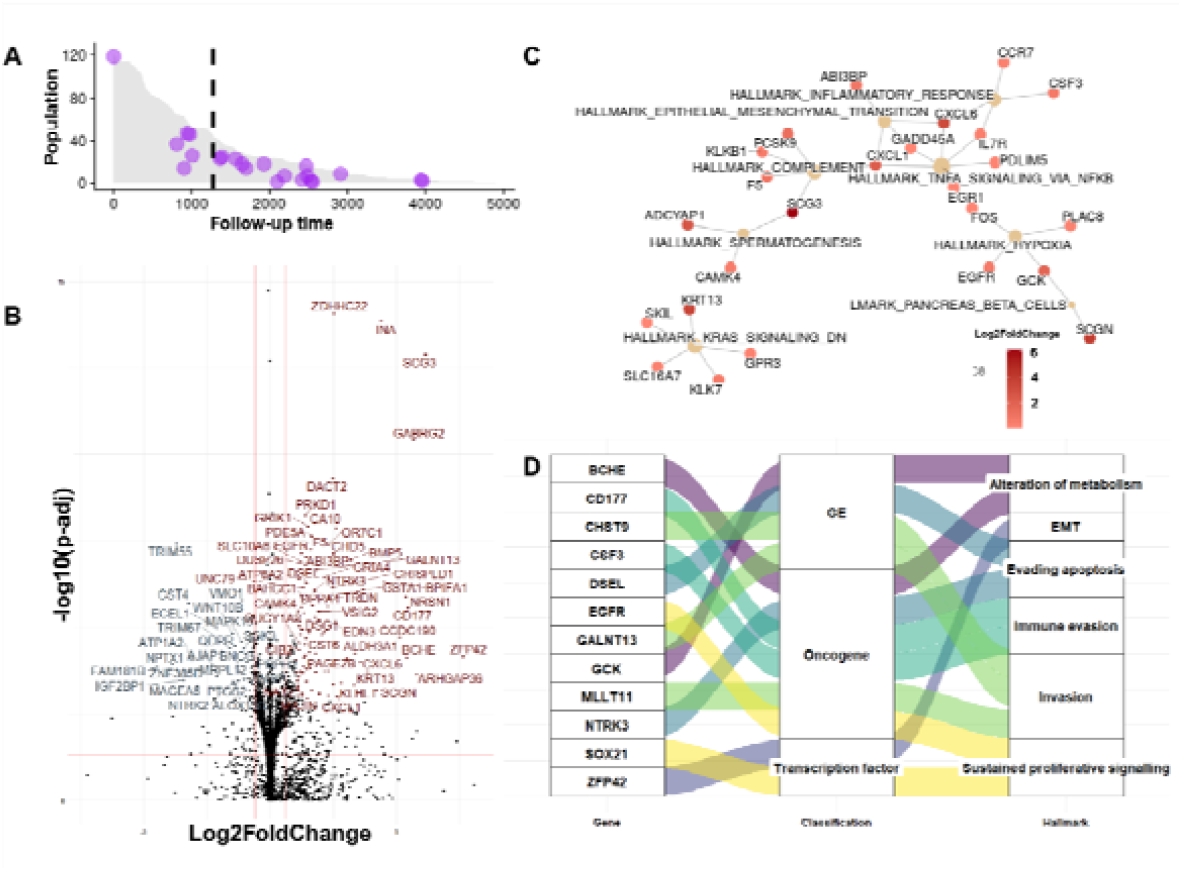
Aberrations in the LumB-HS Late Hazardous group. **(A)** Population-time plot of the LumB-HS group. Dots indicate death events, dotted line at 1,273 days indicates where the hazard ratio surpasses 1. **(B)** Volcano plot showing up- and down-regulated genes in the LumB-HS Late Hazardous patients compared to the LumB-HS Early Protective patients. **(C)** Enrichment of hallmark pathways in the LumB-HS Late Hazardous group. **(D)** Upregulated transcription factors, oncogenes, and GEs activate the hallmarks of cancer.

A differential expression analysis was performed on the two groups to identify potential drivers of tumorigenesis in the Late-Hazardous group. There were 105 upregulated and 24 downregulated genes in the LumB-HS Late-Hazardous group (p-adj < 0.05; Log2FoldChange > 1) (**Figure 4B**). WNT10B, FAM181B and NPTX1 were among the downregulated genes, whereas BMP5 and BPIFA1 were upregulated alongside GEs GALNT13, DSEL, and CHST9. The upregulation of these genes in the LumB-HS Late Hazardous group resulted in enrichment of several tumorigenic pathways such as the inflammatory response, hypoxia, and epithelial to mesenchymal transition **(Figure 4C)**. The simultaneous upregulation of transcription factors such as SOX21 and ZFP42, oncogenes like BCHE, CD177 and EGFR, as well as glycoenzyme genes suggests a highly co-ordinated effort to drive oncogenesis via activation of the hallmarks of cancer. These hallmarks include evasion of the immune system, sustained proliferative signalling, and epithelial to mesenchymal transition **(Figure 4D)**.

## Discussion

Oncogenesis is driven by the consistent proliferation and maintenance of cancer stem cells that hijack embryonic developmental circuitry to achieve this goal. Glycosylation controls the activation of these developmental pathways through modulation of proteins and lipids in the tumour and surrounding matrix.^3^ The extensive heterogeneity of breast cancer suggests that a variety of tumorigenic mechanisms deployed in different ways are responsible for the range of clinically observable outcomes. These mechanisms may well be mediated by the activation of distinct glycosylation pathways in each class of outcome.

To this end, we find that glycosylation signatures within Basal and Luminal B breast cancer reflect a combination of PAM50 subtype and mRNAsi, as elucidated through GE-based unsupervised clustering **(Figure 1A)**. The integration of PAM50 subtype and mRNAsi class labels led to the discovery of clinically useful insights. Patients in the Basal-HS group have improved outcomes as compared to the LumB-HS patients **(Figure 1B)**. An unexpected discovery is that high stemness in the Basal-HS group is a protective factor, while in the LumB-HS group, the risk conferred by high stemness increases gradually over time. Specifically, for LumB-HS patients, high stemness is a hazardous prognostic factor that radically evolves and becomes definitive after 1,273 days from initial diagnosis **(Figure 1C)**. This variation in patient outcome with respect to subtype and stemness goes against the grain of conventional understanding, which associates stemness with increased tumour aggression, metastasis, and recurrence.^62^ This unusual observation is rationally understood in the context of differential glycosylation that underpins the biochemical networks and signalling pathways. The resultant implication from this perspective is that glycosylation opens an avenue to detect the biochemical changes driving the observed differences in clinical outcomes between the subtypes as a result of high stemness.

Using differential gene expression analyses between the LumB-HS and Basal-HS groups the isolation of genes that exhibited subtype- and stemness-specific variation **(Figure 2A)** became possible. The expression of the 184 candidate genes in the Basal-HS group resulted in an upregulation of the immune response, including functions such as T-cell and lymphocyte activation, and leukocyte cell adhesion **(Figure 2B)**. By activating or suppressing the inflammatory response, immune and stem cells work together to mediate the balance between self-renewal and differentiation.^9^ These interactions are highly complex and incompletely understood. While the immune system is critical for the elimination of cancer cells, constant assaults on the immune system lead to immune exhaustion and eventual metastasis.^63^ Despite the potential risks that an active immune response may confer to patients, the pro-inflammatory milieu in Basal-like breast cancer appears to be beneficial,^64^ further evidenced by the positive association between CD274 and IFNG expression, and patient survival in this cohort.

In contrast, the LumB-HS group is characterised by metabolic disturbances that promote tumour aggression, such as activation of oxidative phosphorylation **(Figure 2C)**. Oxidative phosphorylation is critical for stem cell maintenance and enhances the metastatic potential of tumours.^65^ SOX2, a transcription factor that facilitates stem cell self-renewal, promotes tumour dependence on oxidative phosphorylation. Interestingly, SOX2 expression is dependent on an acidic microenvironment,^66^ which is facilitated by the upregulation of ASIC3.^67^ Thus, stemness appears to be maintained via activation of distinct biochemical pathways between the LumB-HS and Basal-HS groups.

Concentrations of metabolites were predicted to vary between the two groups **(Figure 2D)**. The metabolites enriched in the Basal-HS group form part of the surrounding extracellular tumour milieu. Galactose is a structural component of tumour microenvironment (TME) receptors, including immune and stem cells.^68^ Malonyl-CoA, an intermediate metabolite in fatty acid metabolism, is a key effector of immune cell activation, immunosuppression, and TME communication.^69^ The LumB-HS metabolites 4a-Hydroxy-BH4^70^ and Pantetheine-4’-Phosphate^71^ protect cells against oxidative stress, while GDP-Mannose is a precursor to protein mannosylation. An additional difference between these two groups is the increased abundance of M0 and M1 macrophages in the Basal-HS group **(Figure 3E)**. Macrophage polarisation towards an M1 phenotype is typically associated with a pro-inflammatory and anti-tumorigenic microenvironment.^61^ Thus, the functional distinctions between these two groups are reflected in metabolite concentration and immune cell abundance, further explaining phenotypic and clinical variation.

Of the 442 identified candidate genes, 19 are GEs, which distinguished between the two groups with 88.47% accuracy **(Figure 2F)**. GALNT3, an O-GalNAc transferase, is the most important classifier and upregulated in the Basal-HS group, along with HPSE and ARSB, which sulfate proteins and lipids in the extracellular matrix. Additionally, the Basal-HS group exhibits upregulation of HS3ST6, HYAL4, and FUT2, all involved in modification of cell surface glycoconjugates. Glycosylation regulates immune system function through cell surface receptor modification, driving the observed Basal-HS phenotype.

In contrast, important candidate GEs in the LumB-HS group are involved in mannosylation and correct trafficking of proteins through the ER-Golgi complex (POMGNT1, LMAN2, GMPPB, DPY19L4). Although glycosylation and the oxidative stress response are traditionally thought of as independent pathways, evidence suggests highly orchestrated and reciprocal interactions between the two dictate cellular response to microenvironmental shifts.^72^ While the mechanisms are yet to be uncovered, observational studies note that highly mannosylated N-glycans are a key feature of embryonic stem cells^73^ and promote metastasis in cancer.^74^

Candidate GEs between the two groups show unique correlation patterns **(Figure 3A)**, further suggesting that GE upregulation coincides with activation of separate functional pathways. Through functional profiling of correlated genes, distinct roles for glycosylation in tumorigenesis emerged **(Figure 3B)**. Sulfated glycosaminoglycans, synthesised by enzymes such as HPSE and ARSB, modulate the immune response by attracting and facilitating adhesion of immune cells, and binding or sequestering cytokines.^63^ Indeed, HPSE is critical for natural killer activation and immune infiltration,^75^ as well as polarisation of macrophages towards an M1 phenotype.^76^ Furthermore, sulfation of glycosaminoglycans control activation of ERK and Wnt signalling pathways, thereby regulating the balance between stem cell self-renewal and differentiation.^77^ While a pro-inflammatory state may lead to immune exhaustion, GALNT3 prevents immunosuppression through inhibition of NFKB and cMET-Akt.^78^ However, the expression of GALNT3 has been associated with increased tumour aggression,^79,80^ highlighting the context-dependence of the immune system in cancer.

Autophagy, Notch and mTOR signalling were correlated to protein trafficking and mannosylation GEs, as well as to HK1, which facilitates the first step of glycolysis. While glycolysis and oxidative phosphorylation are thought of as mutually exclusive, recent perspectives on the topic suggest that these processes may be coupled in cancer.^81^ Interestingly, HK1 is correlated to ZMIZ1, an oncogene that regulates Notch signalling^82^ and may modulate glucose homeostasis.^83^ Furthermore, dysregulation of autophagy, mTOR signalling and ER stress lead to evasion of apoptosis.^84,85^ Considering the significant crosstalk between the ER, Golgi, and mitochondria, it is not surprising that these biochemical aberrations appear to be highly co-ordinated.

The time-dependent mechanism through which stemness confers increased risk to LumB-HS patients **(Figure 4A)** was elucidated through differential expression analysis. After 1,273 days – around 3.5 years – from initial diagnosis, expression of self-renewal genes WNT10B and FAM181B are downregulated, while metastasis-promoting genes BMP5 and BPIFA1 are upregulated **(Figure 4B)**. Stem cells can be thought of as templates for tumorigenesis, differentiating into specialised cells depending on the needs of the tumour over time. These findings reflect the switch between self-renewal and specialisation that must eventually occur for a tumour to metastasise. Indeed, the upregulation of aggression-promoting genes result in enhanced EMT, a hypoxic microenvironment, and an inflammatory response **(Figure 4C)**.

The inflammatory response is upregulated in the LumB-HS Late Hazardous group, along with immunosuppressive genes such as CD177 and DSEL, which is associated with higher immune infiltration but worse patient outcome.^86^ This contrasts with the observed positive effect of inflammation in the Basal-HS cohort and further emphasises the context-specificity of the immune system in cancer. The increased risk of high stemness in the LumB-HS group likely occurs as a result of highly orchestrated events mediated by the upregulation of key glycogenes, oncogenes, and transcription factors, that drive the hallmarks of cancer **(Figure 4D)**. Indeed, CHST9 is transcriptionally targeted by SOX21 and ZFP42, involved in cellular proliferation^87^ and epigenetic modification^88^ respectively. These results provide further evidence that glycosylation forms part of an organised set of events that drive tumorigenesis.

Approaching cancer through the lens of aberrant glycosylation disentangles the biochemical nuanced networks that drive context-specific variations in phenotype and patient outcome. Analysis of glycosylation signatures within high stem Basal and LumB breast cancer expose shortcomings in the conventional prognostic patterns assumed from these hormonal and stemness markers. Instead the discovery of distinct pathways that are used to maintain the degree of stemness is more telling of patient outcome. These finding show that glycosylation is central to the tumorigenic process, and incorporating these pathways into the mainstream understanding of cancer will invariably lead to novel hypothesis generation and improved diagnostic and therapeutic strategies.

## Authors’ contributions

MT: Conceptualisation, data curation, formal analysis, visualisation, methodology, writing original draft, editing. KJN: Conceptualisation, methodology, funding, supervision, writing.

## Acknowledgements

This work is based in part upon research supported by the National Research Foundation (NRF) CPPR 466624 grant (KJN) and the South African Medical Research Council Self-Initiated Grant (SAMRC SIG) 416090 grant. MT thanks the National Research Foundation (NRF) for a Doctoral Postgraduate Scholarship (PMDS22051711715), as well as the University of Cape Town for a Doctoral Research Scholarship. We thank the Centre for High Performance Computing (CHPC) for computational resources (CHEM0840).

