## Supplementary_Figure_S1 and Supplementary_Table_S1 for "Aberrant glycosylation reveals unexpected clinical outcomes between Luminal B and Basal high stemness index breast cancer cohorts": Supplementary_Figure_S1_GEStemCell.docx

### Supplementary Information

#### Supplementary Figures

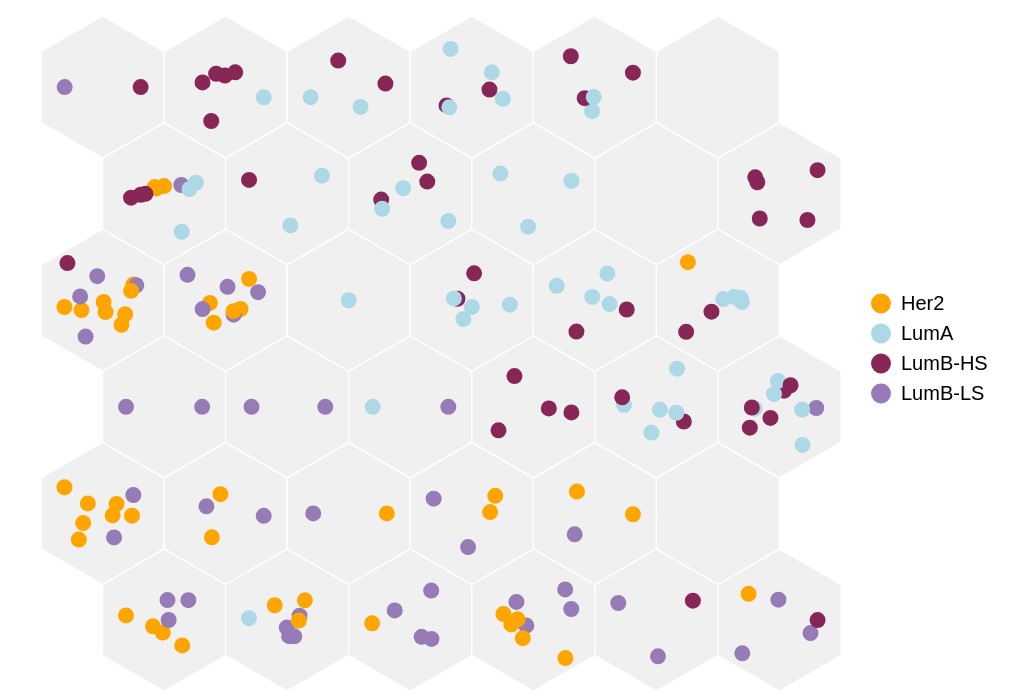

**Supplementary Figure S1:** Glycosylation-based clustering of non-Basal samples highlights the similarity between LumB-LS and HER2 samples.
